# The catscape: spatial manifestation of a pet cat population with outdoor access

**DOI:** 10.1101/2021.12.20.473457

**Authors:** Richard Bischof, Nina Rosita Hansen, Øyvind Skarsgard Nyheim, Astrid Kisen, Lillian Prestmoen, Torbjørn Haugaasen

**Author notes:** **Correspondence** Richard Bischof, Faculty of Life Sciences and Natural Resource Management Norwegian University of Life Sciences, Høgskoleveien 12, Ås, Norway.

## Abstract

The domestic cat (*Felis catus*) is the most popular companion animal and the most abundant carnivore globally. It is also a pet with an immense ecological footprint, because even non-feral and food-subsidized cats are prolific predators. Whereas knowledge about the spatial behavior of individual domestic cats is growing, we still know little about how a local population of free-ranging pet cats occupies the landscape. Using a citizen science approach, we GPS-tagged 92 pet cats with outdoor access living in a residential area in southern Norway. The resulting position data allowed us to construct both individual home range kernels and a population-level utilization distribution. Our results reveal a dense predatory blanket that outdoor cats drape over and beyond the urban landscape. It is this population-level intensity surface - the “catscape” - that potential prey have to navigate. There were almost no gaps in the catscape within our residential study area and therefore few terrestrial refuges from potential cat predation. However, cats spent on average 79% of their outdoor time within 50 meters to their owner’s home, which suggests that the primary impact is local and most acute for wildlife in the vicinity to homes with cats. We discuss the catscape as a conceptual and quantitative tool for better understanding and mitigating the environmental impact of domestic cats.

## Main

Pet cats with outdoor access are as much part of ecological communities as wildlife species that tolerate or benefit from proximity to humans. Despite receiving food subsidies from their human owners, many domestic cats are prolific predators and prey on a wide range of wildlife species [1, 2]. With an estimated 600 million pet cats worldwide [3], this is the most abundant carnivore, wild or domestic. Consequently, the quantity and diversity of wild prey killed by domestic cats is staggering [1, 2, 4].

The direct ecological impact of pet cats is linked with their outdoor activity, and a growing number of studies - often using GPS tracking - seeks to elucidate their space use behavior. Studies range from local [5] to regional [6], including an extensive investigation involving over 900 individual cats from 6 countries [3]. Thanks to such studies, we are beginning to understand the space use patterns of outdoor cats. However, we are not aware of any study that has attempted to track all or the majority of cats within a neighborhood with a typical cat density. This is a surprising gap in information, as the local ecological impact of pet cats is attributable to their sheer numbers [7].

Wildlife preyed upon by cats face the combined spatial representation of the local cat population, which in urban areas readily reaches densities of several hundred cats per km^2^ [8–10]. Mapping the “catscape” - the combined intensity of space use by a local cat population - could help us better understand and mitigate the ecological impacts of domestic cats. During a month-long study, and with the help of citizen scientists, we GPS-tagged and tracked 92 domestic cats living with their owners in a small (1 km^2^) residential area in southern Norway. This unprecedented density of GPS-tagged cats (87/km^2^) and the analytical approach we used resulted in an ecologically meaningful representation of a cat population - spatially-explicit risk from a prey’s perspective.

## Results & Discussion

### The Catscape

Our analysis revealed that individual utilization distributions (UDs, Fig. 2) of pet cats with outdoor access coalesce into a joint surface that drapes over the suburban landscape and the surrounding areas (Fig. 3). It is this surface that potential prey species have to navigate, and its characteristics not only convey the spatial configuration of risk, but can also guide the selection and targeted application of mitigation measures.

**Figure 1:**
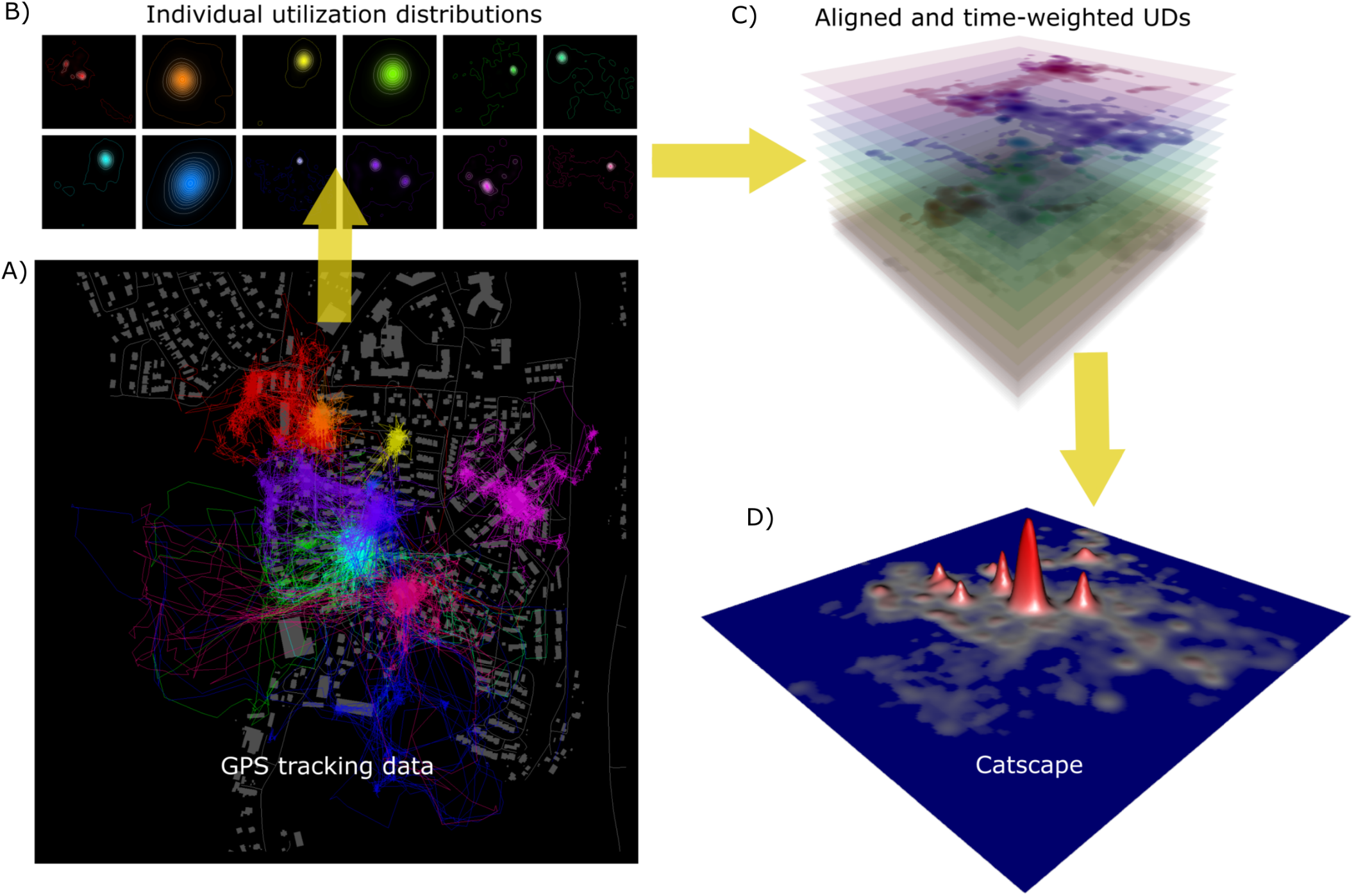
Illustration of the construction of the catscape by aggregating the utilization distributions (UDs) of 12 example pet cats. High-throughput GPS data (A) are used to estimate individual UDs with Brownian bridge motion models (B). Individual UDs are weighted according to the average time spent outdoors on days with data (C), and are then summed across individuals to yield the combined intensity of use (D).

**Figure 2:**
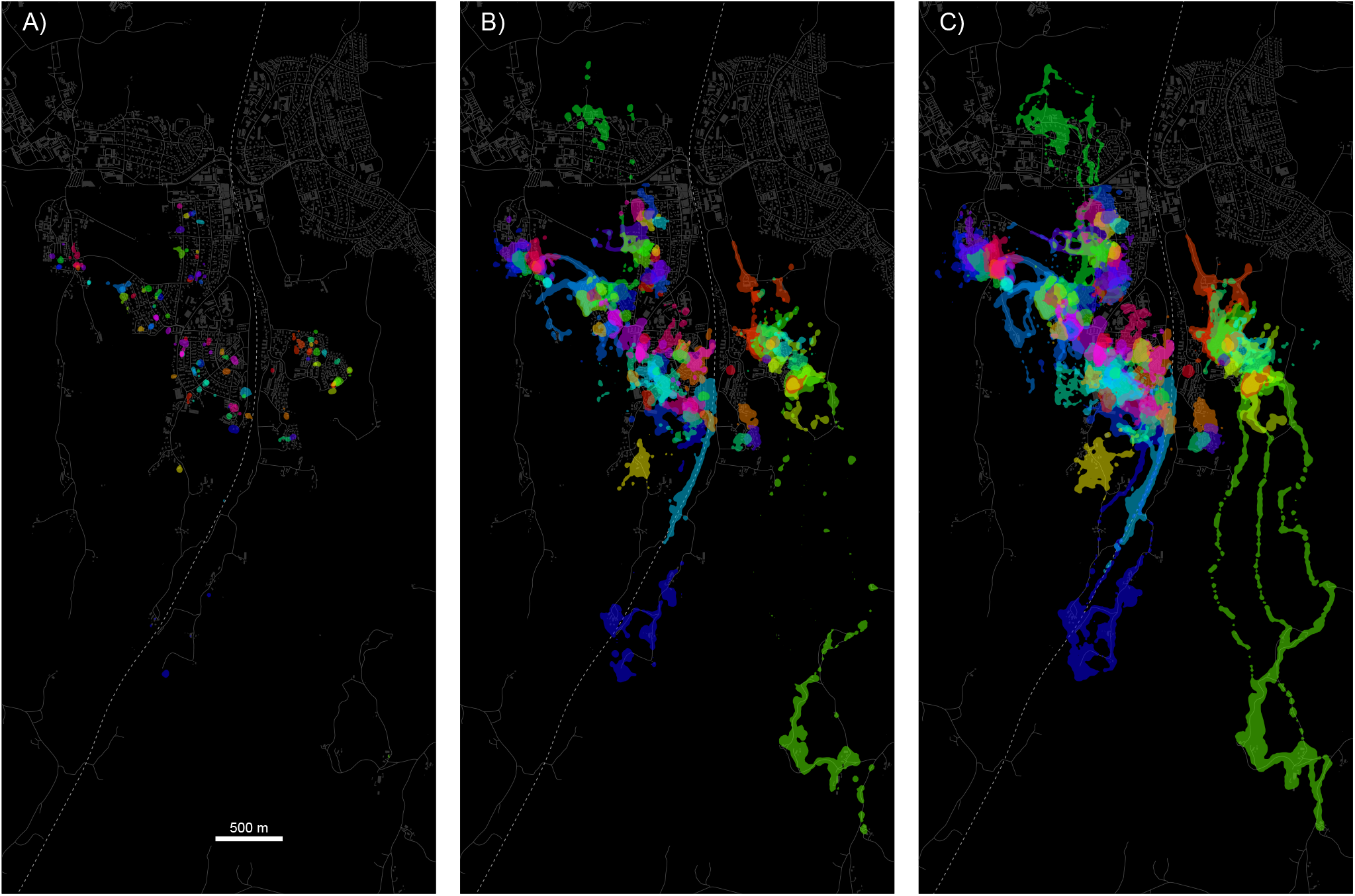
Individual home ranges for 92 pet cats (color-coded) that were GPS-tracked for up to one month in a residential area in southern Norway. Shown are 50% (A), 95% (B), and 99% (C) home ranges, based on corresponding vertices of Brownian bridge motion models. Cats spent the majority of their outdoor time in close proximity to their owner’s home, evident in the small core areas (50% home range) compared with the 95% and 99% home range polygons. Grey lines and polygons in the background indicate roads and buildings, respectively. A fenced railroad (dashed line) transects the study area.

**Figure 3:**
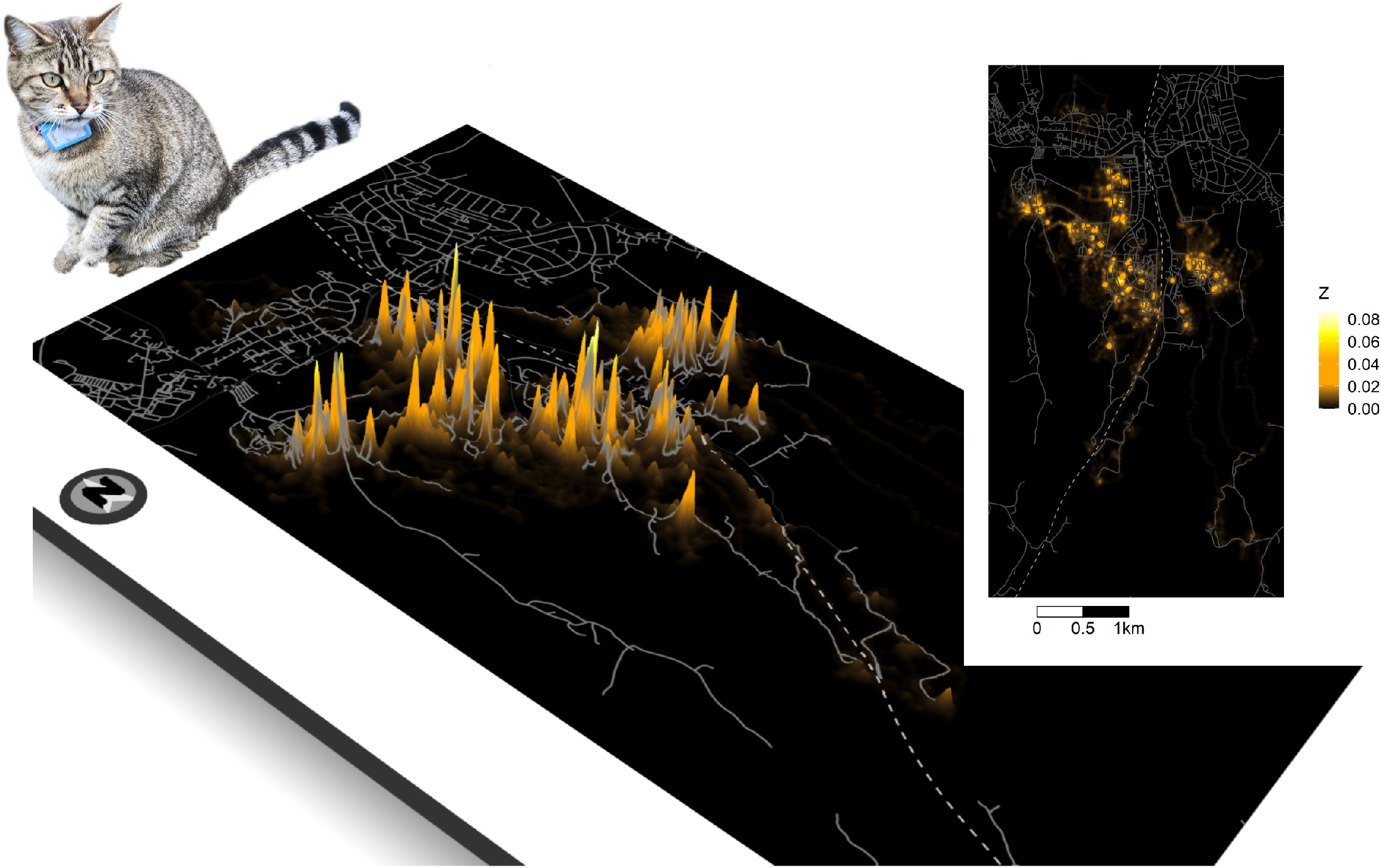
Three-dimensional representation of the catscape (with 2D inset), constructed using individual utilization distributions of 92 domestic cats. The height of the surface (*Z*, square-root transformed for visualization) denotes the intensity of use by cats. Grey lines indicate roads; a fenced railroad (dashed line) transects the study area.

The catscape represents the combined utilization intensity of all GPS-tracked cats. The height of the surface is thus a function of both the number of cats that use a particular location and the amount of time they spend there. The surface is highly variable; within the region delineated by the 99% vertex of the catscape, we measured a 1660-fold difference between the highest and lowest utilization intensity values (Fig. 3). The catscape shown here represents a minimum use-scenario. Based on known cat owners that chose not to participate and ancillary information from 47 camera traps distributed throughout residential backyards in the study area (Extended Data Fig. S1), we estimate that we tracked at least 73% of the cats living inside the study area and approximately 40% of the outdoor activity by all cats. Activity missed is likely a combination of activity by participant cats during times when they were not wearing a functioning collar, activity by non-participant cats that live inside the study area, and activity by cats entering the study area from outside.

Although seemingly high, the number of known cats (126, density: 119/km^2^) with homes in our study area is comparable or lower than values reported elsewhere. For example, domestic cat densities in urban areas in the UK have been estimated to range between 131.8 and 1 579,2 cats/km^2^ (median 417.3 cats/km^2^) [8] and average cat densities of above 200 per km^2^ have been reported by studies in New Zealand [9] and the US [10]. In Norway, with a human population size of 5.4 million, there are an estimated 770 000 pet cats [11].

One approach to mitigate environmental impacts of cats is to decrease the spatial coverage and overall height of the catscape. This could be accomplished by reducing the number of cats with outdoor access, including prohibiting cat ownership in sensitive areas [12], or limiting the amount of time cats spend outdoors [13]. Reduction in pet cat numbers, while theoretically feasible, presumably involves the kind of political and regulatory maneuvering most jurisdictions shy away from [14]. Moreover, without a better understanding of the role of intra-species interactions in the spatio-temporal dynamics of cat populations, it is unclear whether the reduction of cat numbers or the duration and frequency of outdoor access would lead to a corresponding reduction in overall space utilization [12]. It has been suggested that removing cats in a dense population may lead to a reduction in predation pressure, but not in a reduction of spatial coverage, as remaining cats expand their home ranges due to density dependent effects on movements [9]. Counter-intuitive effects of predator control on predation have been demonstrated or suggested also in other species, for example due to the disruption of social structure, changes in behavior of remaining individuals, or knock-on effects on other predator species [15].

The spatial coverage of the catscape can be reduced by shrinking the average home range size of constituent cats. Confinement [13], including curfews to keep cats indoors during times with more pronounced roaming behavior [5], and sterilization [16, 17] have been suggested to reduce roaming behavior and home range size in domestic cats. This, like a reduction in the number of cats, would decrease the total size of the area impacted and create gaps in the catscape that could serve as refuges for potential prey.

### Beyond urban

Thirty-eight percent (0.61 km^2^) of the catscape (1.59 km^2^ based on the area delineated by the 99% volume contour of the catscape) extended beyond urban areas and areas with infrastructure (roads, farm yards, etc.). That domestic cats living in urban areas range into natural areas has already been demonstrated [3, 18]. Our study revealed the magnitude of this halo at the population-level and the highly variable contribution that individuals make to it, with 10% of participant cats accounting for 63% of the non-urban habitat use. However, because cats in this study lived primarily in urban neighborhoods and individual space utilization declines rapidly from each cat’s core use area (Fig. 2), the majority (90%) of the risk represented by the catscape fell within urban habitat.

Methods that decrease cat density and home range size/ranging behavior would potentially also reduce the extent to which the catscape and associated impacts bleed into non-urban areas. Another strategy could be to tolerate medium and large carnivores around urban areas, as these may provide an added ecological service through culling of roaming cats or impacting cat space use behavior [19–22] by creating a landscape of fear [23, 24].

### Spiked utilization

The catscape is characterized by pronounced peaks, reflecting high site fidelity of cats to their owners’ homes, with further clustering evident in both individual and population-level UDs (Fig. 3). At the population-level, cat utilization intensity was 9.5-fold higher within 50 m of a cat owner’s home than at a distance of 50 m to 100 m. Similarly, at 0.2 ha (95% CI: 0.1 - 0.5 ha; Fig. 2A), the core area of the average cat (50% UD; usually centered on the owner’s residence) was less than 1/14th the size of its 95% home range (mean: 2.6 ha; 95% CI: 0.3 - 11.2 ha; Fig. 2B).

Concentrated activity characterizes the spatial behavior of most pet cats [3] and suggests that the highest impact is local and most acute for wildlife utilizing space in the vicinity of residences with cats. The configuration of the catscape into pronounced hot-spots offers opportunities for more targeted mitigation measures. This could involve the removal or enhanced protection of bird feeders, nest boxes, and bird baths near the homes and in the yards of cat owners. Here too, un-intended consequences are possible if, for example, the reduction in predation opportunities leads to an expansion of the cat’s activity area, with risk simply shifting farther away from the cat’s home.

### Individual variation

Individual variation in space use was high among cats in this localized population. For example, home range size estimates during the study period ranged from 0.3 to 22.1 ha (mean: 2.6 ha), the extent to which home ranges included non-urban areas ranged from 0% to 66% (mean 10%), and individual cats spent between 8% and 100% (mean 80%) of their time within 50 m of their owner’s home. High intraspecific variation in cat space use behavior has been reported by others [3, 25] and likely propagates to variation in environmental impacts. The high amount of individual variation in space use and other behaviors poses a critical obstacle for the common approach in wildlife ecology of scaling inferences to the population-level from a limited sample of individuals. To be representative, the sample would have to capture the population-level breadth and distribution of the parameter of interest, which is difficult to achieve and verify [26]. Furthermore, investigations targeting individual interactions or the manifestations thereof (e.g., territory configuration and turn-over) are impeded when the majority of individuals remains hidden from the observer; the behavior of tagged individuals is interpreted with the sample in mind, whereas in reality it is influenced by additional, unobserved animals.

Substantial individual variation in space use may also mean that mitigation efforts are more effective if they are customized based on individual characteristics. This is plausible, as inexpensive, effective, and accessible pet tracking devices now make it possible for most owners to remotely monitor the outdoor movements of their pets. This offers opportunities for cat-specific mitigation measures that owners can implement based on information about their cat’s movements and in relation to specific husbandry practices. At a minimum, owners could employ selective containment of cats that exhibit roaming behavior. Owners could also experiment with and adopt maintenance regimes (diet, behavioral enrichment [27]) that reduce roaming behavior of individual cats, even if the impact of these as general interventions may be questionable [25].

### Scaling-up monitoring through citizen science

The individual organisms and their impacts scale up to the manifestation of populations. The same is not always true for ecological studies; information based on a small subset of individuals (or individuals originating from a patchwork of sites) are notoriously unreliable for scaling up to population-level inferences [28, 29]. In our study, the synchronous collection of spatial data on a large proportion of cats in one area was achieved through the participation of the owners of these cats, that is, citizen scientists. Data collected at or near the population scale can help answer important questions that remain elusive to studies with small samples or samples scattered over vast areas. Examples of relevant topics concerning domestic cats are the identification of determinants of space use and movement that take into account intraspecific interactions and, eventually, measuring the link between space use and predation pressure exerted. A particularly important line of inquiry concerns the systematic assessment of the impact of mitigation measures at the population scale.

Citizen science can be an efficient and cost effective approach for collecting data at scales that are otherwise difficult to attain. The associated limitations in control and inherent biases may to some extent be assessed and accounted for by accompanying independent survey methods, such as camera trapping in our study. Citizen science can also serve as a vehicle for motivating, informing, and evaluating measures for mitigating the ecological impacts of cats, as most interventions will need to be implemented by the cat owners themselves. We agree with [14], given the multitude of circumstances faced and attitudes exhibited by cat owners, multidimensional and customizable mitigation strategies are a more likely path to conservation success than general policies.

## Methods

### Study area

The study took place in and around Ås, a small (10 725 inhabitants; 4.73 km^2^) university town in southern Norway made up of campus and a commercial center surrounded by residential areas. The area from which cats were recruited encompassed 1 km^2^ (Fig. 2, Fig. S1) of residential area (primarily single and multi-family homes and yards). A fenced railroad dissects the study area (Fig. 2), with the two sides connected by pedestrian underpasses. The study area is surrounded by a mixture of forest, and agricultural fields, with a moderately undulating topography.

### Cat recruitment

We used several methods to recruit as many cat owners in our study area as possible: flyers distributed in every residential mailbox in the study area, an advertisement on a local social-media group, and word-of-mouth. This redundancy in outreach was motivated by the goal to recruit as many of the resident cats as possible. We also identified potential cat owners that had not registered during the initial recruitment campaign by inquiring about other households with cats in registered participants’ neighborhoods. These additional households with cats were approached directly with an invitation to participate.

Participants registered for the study through an online registration form, which in addition served as a questionnaire for collecting basic information about the cat(s) (e.g., number of cats with outdoor access) and relevant features of the household (e.g., the owner’s contact information). Once registered, participants completed a follow-up online questionnaire collecting detailed information about each cat and its maintenance.

### GPS tracking

GPS tracking took place between May 1 and May 29, 2021. Owners were instructed to affix the collar with the GPS unit to their cat each time the cat exited the home. We used i-gotU GT-120 GPS units (Mobile Action Technology, Inc., Taiwan) weighing 26 g and requiring manual download. GPS units were set to attempt one position fix every 30 seconds while on. Such high-throughput telemetry data are becoming increasingly common and are particularly useful for revealing detailed movement patterns [30]. Data were stored on-board and downloaded by the authors at least once during the tracking period (to ensure proper functioning, rectify errors in tagging/data collection, backup GPS data) and at the end of it. Participating cat owners were also provided with a hotline for addressing technical problems and to replace lost GPS units.

### Camera trapping

We placed one wildlife camera trap with infrared flash in the yards of 47 participating cat owners. The camera models we used were Browning Dark Ops HD Pro Trail Camera BTC-6HDP (37), Browning BTC-6HDPX Dark Ops HD Pro (8) and Browning Spec Ops Full HD (2). Cameras were set 0.5 - 1 m above the ground and in places that would maximize the probability of recording wildlife and domestic cats entering or moving across the yard, while also protecting the privacy of neighbours. Cameras were programmed to record 10 s videos each time they were triggered. Cameras were placed roughly one week after the GPS tracking started, and remained in the gardens for five weeks. Due to memory limitations, cameras were operational for an average of 22.6 days (sd 12.4). For each cat video, we recorded if the cat wore a GPS collar or not, in order to gauge GPS-coverage of cat activity in the study area during GPS tracking.

### Data analysis

We used R v 4.1.1 [31] for the data processing and the analysis outlined below.

### GPS data pre-processing

GPS data from each cat were subjected to the following sequence of pre-processing steps, based partially on recommendations by [32] and [33] :

1. Removed positions with an elevation outside the range 0 - 300m.
2. Removed positions obtained during the first 2 days of tracking.
3. Removed positions obtained on days where the GPS was picked up for data download.
4. Removed positions with an estimated horizontal position error (EPHA) >= 5000.

### Delineation of outdoor activity

Although owners were instructed to remove and switch off the GPS collar when the cat was indoors, we know from previous surveys (unpublished data) that this was not done reliably in every household, especially for cats that exited and entered the house freely through a cat flap or similar. GPS data revealed a high degree of clustering, particularly in the vicinity of the cat’s home. We used R package GPSeqClus [34] to identify sequential position clusters based on a reported gps error <10 m [33]. As we were interested in outdoor activity, we identified GPS clusters with centroids (see previous subsection) that fell within the spatial extent of the owner’s residence (building spatial data obtained from the Norwegian Mapping Authority) and then removed all relocations associated with these clusters. This ensures that positions used during subsequent analyses can be considered arising from a cat’s outdoor activity.

### Individual utilization distributions

We used R package BBMM [35] to construct a Brownian bridge movement model [36, 37] for each cat in the study. We chose BBMM, as it is better equipped to deal with auto-correlated and clustered position data than the kernel density home range estimator commonly used in wildlife studies [38]. Specifically, BBMM allows estimation of probabilistic space use, while accounting for position error and the uncertainty about paths taken between consecutive positions [37].

### Population-level utilization distribution

With some modification, we used the approach outlined by [37], and subsequently followed by [39] and [40], to construct the population-level utilization distribution (UD) from individual Uds (BBMM). This involved the following steps (see also Fig. 1):

1. Resample individual UD rasters to align (same spatial extent and resolution).
2. Weigh cell values in individual UD rasters with the average proportion of a day the cat was tracked outdoors on days with outdoor position data. This assumes that a) cats were tracked for the entire time they spent outdoors and b) on days without any GPS tracking data, cats spent an average amount of time outdoors, but were not tracked. We know from simultaneous camera trapping data that the former assumption was violated, whereas the latter assumption likely held, based on communications with owners. In general, this means that outdoor utilization was underestimated and that the final population-level catscape represents a minimum use-scenario.
3. Aggregate individual UD rasters into a population-level catscape by summing cell values across the stack of 92 weighted individual UD rasters [37]. Unlike [37], we did not perform further scaling of the resulting raster to sum to 1.

This approach results in an ecologically meaningful representation of a cat population - spatially-explicit risk from a prey’s perspective - that does not require further re-scaling. The approach also allows comparison of population-level UDs within (e.g., day vs. night) and between populations in terms of both their magnitude and shape.

## Acknowledgements

We thank participating owners for making their pets available for this study and performing the tasks necessary for data collection. We also thank F. Sarfi, W. Fan, B. Bachman, G.S. Leikanger, C. Glosli, and R. Steen for help with earlier bouts of cat monitoring and assistance with developing the protocols and methods that we built upon in this study. We thank M. Wonderland for assistance with field work and processing of camera trap data. The study was funded by the Research Council of Norway (projects 283741 and 286886). We thank A. Semper-Pascual, J. Swenson, and A. Vallejo for comments on the manuscript.

## Author contributions

RB and TH conceived the idea and designed the study, with input from AK and NRH. RB, TH, and NRH coordinated the data collection. NRH, ØSN, and LP conducted the field work and collected the data. ØSN and LP processed the camera trap data. RB conducted the analysis and wrote the first draft of the manuscript. All authors contributed to subsequent drafts and gave final approval for publication.

## Additional information

Supplementary Information is available for this paper.

Correspondence and requests for materials should be addressed to R. Bischof.

## References

[1] Seymour, C. L. et al. Caught on camera: The impacts of urban domestic cats on wild prey in an African city and neighbouring protected areas. Global Ecology and Conservation 23, e01198 (2020).

[2] Mori, E. et al. License to Kill? Domestic Cats Affect a Wide Range of Native Fauna in a Highly Biodiverse Mediterranean Country 2019.

[3] Kays, R. et al. The small home ranges and large local ecological impacts of pet cats. Animal Conservation 23 (2020).

[4] Loss, S. R., Will, T. & Marra, P. P. The impact of free-ranging domestic cats on wildlife of the United States. Nature Communications 4, 1396 (2013).

[5] Barratt, D. G. Home range size, habitat utilisation and movement patterns of suburban and farm cats Felis catus. Ecography 20, 271–280 (1997).

[6] Moseby, K. E., Stott, J & Crisp, H. Movement patterns of feral predators in an arid environment - implications for control through poison baiting. English. Wildlife Research 36, 422–435 (2009).

[7] Trouwborst, A., McCormack, P. C. & Martínez Camacho, E. Domestic cats and their impacts on biodiversity: A blind spot in the application of nature conservation law. People and Nature 2, 235–250 (2020).

[8] Sims, V., Evans, K. L., Newson, S. E., Tratalos, J. A. & Gaston, K. J. Avian assemblage structure and domestic cat densities in urban environments. Diversity and Distributions 14, 387–399 (2008).

[9] Van Heezik, Y., Smyth, A., Adams, A. & Gordon, J. Do domestic cats impose an unsustainable harvest on urban bird populations? Biological Conservation 143, 121–130 (2010).

[10] Lepczyk, C. A., Mertig, A. G. & Liu, J. Landowners and cat predation across rural-to-urban landscapes. Biological Conservation 115, 191–201 (2004).

[11] Heggøy, O. & Shimmings, P. Huskattens predasjon på fugler i Norge. En vurdering basert på en litteraturgjennomgang tech. rep. (2018), 36.

[12] Morgan, S et al. Urban cat (Felis catus) movement and predation activity associated with a wetland reserve in New Zealand. Wildlife Research - WILDLIFE RES 36 (2009).

[13] Calver, M., Grayson, J., Lilith, M. & Dickman, C. Applying the precautionary principle to the issue of impacts by pet cats on urban wildlife. Biological Conservation 144, 1895–1901 (2011).

[14] Crowley, S., Cecchetti, M. & Mcdonald, R. Diverse perspectives of cat owners indicate barriers to and opportunities for managing cat predation of wildlife. Frontiers in Ecology and the Environment 18 (2020).

[15] Treves, A., Krofel, M., Ohrens, O. & van Eeden, L. M. Predator control needs a standard of unbiased randomized experiments with cross-over design 2019.

[16] Castañeda, I. et al. Trophic patterns and home-range size of two generalist urban carnivores: a review. Journal of Zoology 307 (2018).

[17] Ferreira, G. A., Machado, J. C., Nakano-Oliveira, E., Andriolo, A. & Genaro, G. The effect of castration on home range size and activity patterns of domestic cats living in a natural area in a protected area on a Brazilian island. Applied Animal Behaviour Science 230 (2020).

[18] López-Jara, M. J. et al. Free-roaming domestic cats near conservation areas in Chile: Spatial movements, human care and risks for wildlife. Perspectives in Ecology and Conservation (2021).

[19] Kennedy, M., Phillips, B. E. N. L., Legge, S., Murphy, S. A. & Faulkner, R. A. Do dingoes suppress the activity of feral cats in northern Australia? Austral Ecology 37, 134–139 (2012).

[20] Crooks, K. R. & Soule, M. E. Mesopredator release and avifaunal extinctions in a fragmented system. English. Nature 400, 563–566 (1999).

[21] Ferreira, J. P., Leitão, I., Santos-Reis, M. & Revilla, E. Human-related factors regulate the spatial ecology of domestic cats in sensitive areas for conservation. PLOS ONE 6, e25970 (2011).

[22] Brook, L. A., Johnson, C. N. & Ritchie, E. G. Effects of predator control on behaviour of an apex predator and indirect consequences for mesopredator suppression. Journal of Applied Ecology 49, 1278–1286 (2012).

[23] Laundre, J. W., Hernandez, L & Altendorf, K. B. Wolves, elk, and bison: reestablishing the “landscape of fear” in Yellowstone National Park, USA. English. Canadian Journal of Zoology-Revue Canadienne De Zoologie 79, 1401–1409 (2001).

[24] Ritchie, E. G. & Johnson, C. N. Predator interactions, mesopredator release and biodiversity conservation. English. Ecology Letters 12, 982–998 (2009).

[25] Hall, C. M. et al. Factors determining the home ranges of pet cats: A meta-analysis. Biological Conservation 203, 313–320 (2016).

[26] Milleret, C. et al. GPS collars have an apparent positive effect on the survival of a large carnivore. Biology Letters 17 (2021).

[27] Cecchetti, M., Crowley, S. L., Goodwin, C. E. D. & McDonald, R. A. Provision of high meat content food and object play reduce predation of wild animals by domestic cats Felis catus. Current Biology 31, 1107–1111.e5 (2021).

[28] Hebblewhite, M. & Haydon, D. T. Distinguishing technology from biology: a critical review of the use of GPS telemetry data in ecology. Philosophical Transactions of the Royal Society B: Biological Sciences 365, 2303–2312 (2010).

[29] Allen, A. M. et al. Scaling up movements: from individual space use to population patterns. Ecosphere 7, e01524 (2016).

[30] Bischof, R., Gjevestad, J. G. O., Ordiz, A., Eldegard, K. & Milleret, C. High frequency GPS bursts and path-level analysis reveal linear feature tracking by red foxes. Scientific Reports 9, 8849 (2019).

[31] R Core Team. R: A Language and Environment for Statistical Computing R Foundation for Statistical Computing (Vienna, Austria, 2021).

[32] Gupte, P. R. et al. A guide to pre-processing high-throughput animal tracking data. bioRxiv, 2020.12.15.422876 (2021).

[33] Morris, G. & Conner, L. Assessment of accuracy, fix success rate, and use of estimated horizontal position error (EHPE) to filter inaccurate data collected by a common commercially available GPS logger. PLOS ONE 12, e0189020 (2017).

[34] Clapp, J. G., Holbrook, J. D. & Thompson, D. J. GPSeqClus: An R package for sequential clustering of animal location data for model building, model application and field site investigations. Methods in Ecology and Evolution 12, 787–793 (2021).

[35] Nielson, M. R., Sawyer, H. & McDonald, T. L. BBMM: Brownian bridge movement model R package version 3.0 (2013).

[36] Horne, J. S., Garton, E. O., Krone, S. M. & Lewis, J. S. Analyzing animal movements using Brownian bridges. Ecology 88, 2354–2363 (2007).

[37] Sawyer, H, Kauffman, M. J., Nielson, R. M. & Horne, J. S. Identifying and prioritizing ungulate migration routes for landscape-level conservation. Ecological Applications 19, 2016–2025 (2009).

[38] Fischer, J. W., Walter, W. D. & Avery, M. L. Brownian Bridge Movement Models to Characterize Birds’ Home Ranges. The Condor 115, 298–305 (2013).

[39] Seidler, R., Long, R., Berger, J., Bergen, S. & Beckmann, J. Identifying impediments to long-distance mammal migrations. Conservation biology : the journal of the Society for Conservation Biology 29 (2014).

[40] Collins, G. Seasonal distribution and routes of pronghorn in the Northern Great Basin. Western North American Naturalist 76, 101–112 (2016).

