## Supplementary information for "The catscape: spatial manifestation of a pet cat population with outdoor access"

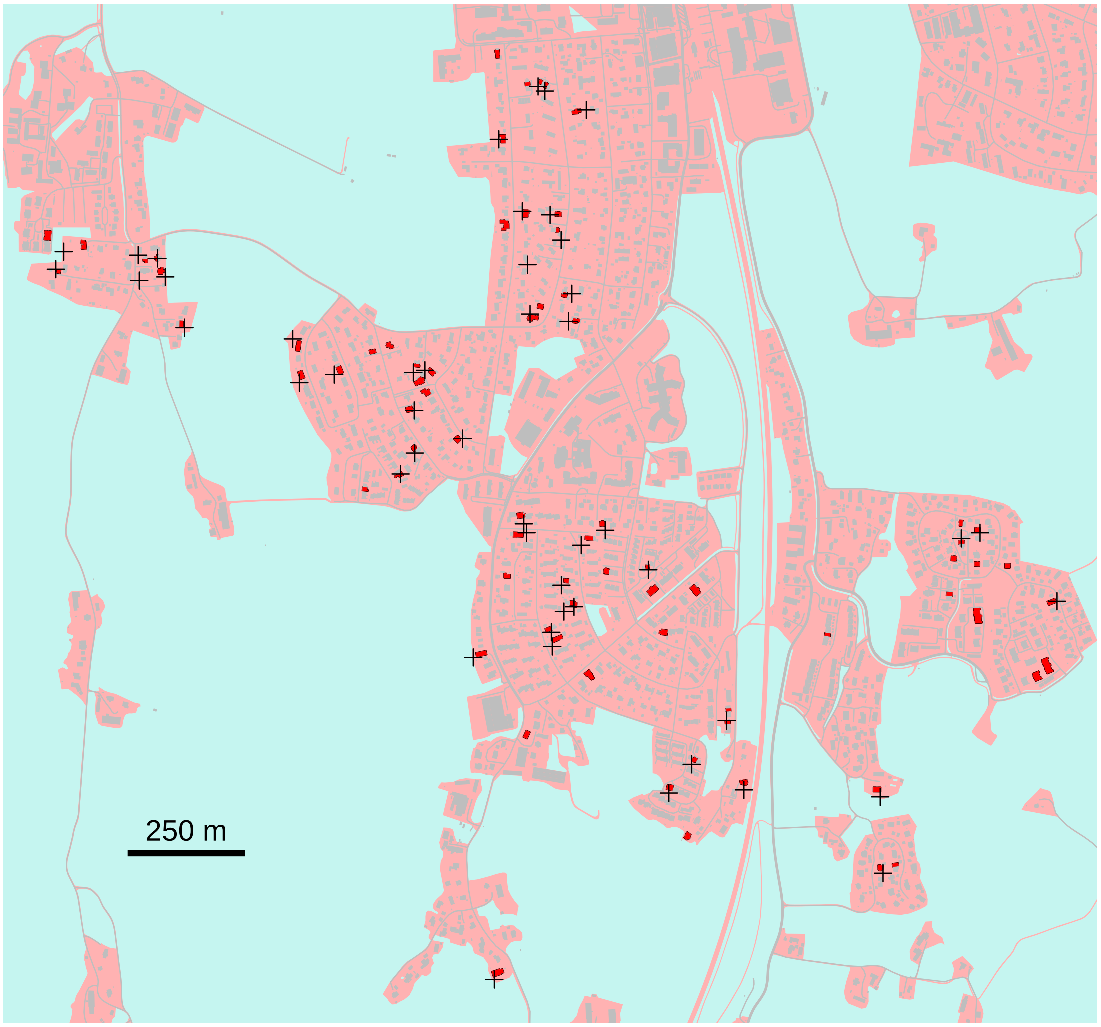


Fig. S 1: Map of the study area showing the distribution of 47 camera traps (black crosses) used to monitor domestic cats within the study area. Camera trap locations are shown in the context of developed areas, including roads (pink), pastures, fields, and forests (turquoise). Buildings are shown in grey, with the homes of 92 GPS-tagged cats in red.
